# Transcriptomic profiling of murine GnRH neurons reveals developmental trajectories linked to human reproduction

**DOI:** 10.1101/2023.06.22.546062

**Authors:** Yassine Zouaghi, Daniel Alpern, Vincent Gardeux, Julie Russeil, Bart Deplancke, Federico Santoni, Nelly Pitteloud, Andrea Messina

## Abstract

Gonadotropin-releasing hormone (GnRH) neurons play a crucial role in human reproduction and are associated with a spectrum of conditions. However, the underlying biological mechanisms remain elusive due to their small number and sparse distribution. We performed transcriptomic profiling of GnRH neurons during mouse embryonic development, revealing their molecular identity and gene expression dynamics. Our findings show that GnRH neurons undergo a profound transcriptional shift as they migrate from the nose to the brain and that distinct expression trajectories are associated with critical biological processes, including cell migration, neuronal projections, and synapse formation. Cell-to-cell communication analysis revealed timely and spatially restricted modulation of signaling pathways involving known molecules, such as Semaphorins and Plexins, and novel candidates, such as Neurexins and Endothelins. Using GWAS genes linked to human reproductive onset, we found a specific association with GnRH neuron trajectories rising in late developmental stages and involved in neuron maturation and connectivity. Finally, analysis of the genetic burden in a large cohort of patients with congenital GnRH deficiency revealed specific GnRH neuron trajectories with a significant mutation load compared to controls.

In conclusion, this study revealed the gene expression dynamics underlying GnRH neuron embryonic development and provides novel insights linking GnRH neuron biology to human reproduction.

## Introduction

Reproductive health encompasses diverse conditions with significant implications for fertility, metabolism, well-being, and aging.^1^ Central to human reproduction are gonadotropin-releasing hormone (GnRH) neurons, which govern the activity of the hypothalamic-pituitary-gonadal (HPG) axis and, ultimately, the secretion of sex steroids.^2^ After their identification in the 1970s, it was found that GnRH neurons originate from the nasal placode and migrate to the hypothalamus during embryonic development.^3-6^ This peculiar ontogenetic event is conserved in humans and, when defective, leads to Kallmann Syndrome, which is characterized by the lack of puberty (congenital hypogonadotropic hypogonadism, CHH) and the sense of smell.^7,8^ Since then, other defects in GnRH neuron biology have been implicated in several reproductive disorders, including normosmic CHH (nCHH), altered timing of puberty, and polycystic ovary syndrome.^9^ However, the genetic and molecular mechanisms linking GnRH neuron biology to these disorders remain unclear.

Despite some inter-species differences, the mouse model is a reference for GnRH neuron migration and has been extremely useful in defining three main developmental stages.^10,11^ (i) At E10.5, GnRH neurons originate in the olfactory placode and migrate as a continuum stream through the nasal mesenchyme and along the olfactory-vomeronasal nerves (ON/VN) and terminal nerve (TN) bundles toward the olfactory bulb (OB).**^4,6,10-12^** (ii) Next, they cross the nasal-forebrain junction (cribriform plate) at the olfactory bulb level, adapt to the new environment and continue their journey following the TN, which turns ventrally toward the hypothalamus. (iii) Once scattered through the hypothalamus, GnRH neurons stop moving and start building their network by connecting with hypothalamic and extra hypothalamic targets, including the median eminence of the hypothalamus (i.e., where the neurosecretion takes place).

The molecular characterization of GnRH neurons has improved dramatically in the past decades, mainly through the extensive use of mouse genetics and precious insight from human genetics.**^12-15^** However, due to their small number and peculiar spatial distribution, GnRH neurons’ precise gene expression dynamics and their drivers remain elusive. Herein, we combined a validated FACS-based method to isolate GnRH neurons with high-throughput RNA sequencing and bioinformatics to map the gene expression changes of GnRH neurons during mouse embryonic development. Further, we investigate how these gene expression dynamics affect GnRH neuron biology and the potential implications for human reproduction.

## Results

### GnRH neurons experience a transcriptional shift from nose to brain

To depict GnRH neuron expression dynamics, we used a low-input-optimized RNA sequencing approach on FACS-isolated GFP^+^ and GFP^-^ cells from *GnRH::GFP* transgenic embryos at four key spacetime points (Fig. 1a). *1) E12.5 nose*: GnRH neurons are almost exclusively in the nose while moving caudally toward the brain. *2) E14.5 nose*: about half of the GnRH neurons are still migrating throughout the nose. *3) E14.5 brain*: the remaining GnRH neurons have already entered the brain and turned ventrally toward the hypothalamus. *4) E18.5 brain*: virtually all GnRH neurons reached their final location in the brain and have completed their journey. The identity of the sorted cells was confirmed *in silico* by measuring *Gnrh1* transcript relative to GFP expression. Notably, *Gnrh1* transcript was expressed within GFP negative samples, albeit in a low range (Fig. 1b), indicating that a small fraction of GnRH neurons might not express the GFP reporter, as previously shown in transgenic reporters.^16,17^ The *bonafide* location of GnRH neurons (i.e., microdissection specificity) was confirmed by the overlap of the most differentially expressed genes in all GFP negative samples with specific markers of developing nose and brain tissues (Fig. 1c, d).

**Figure 1.**
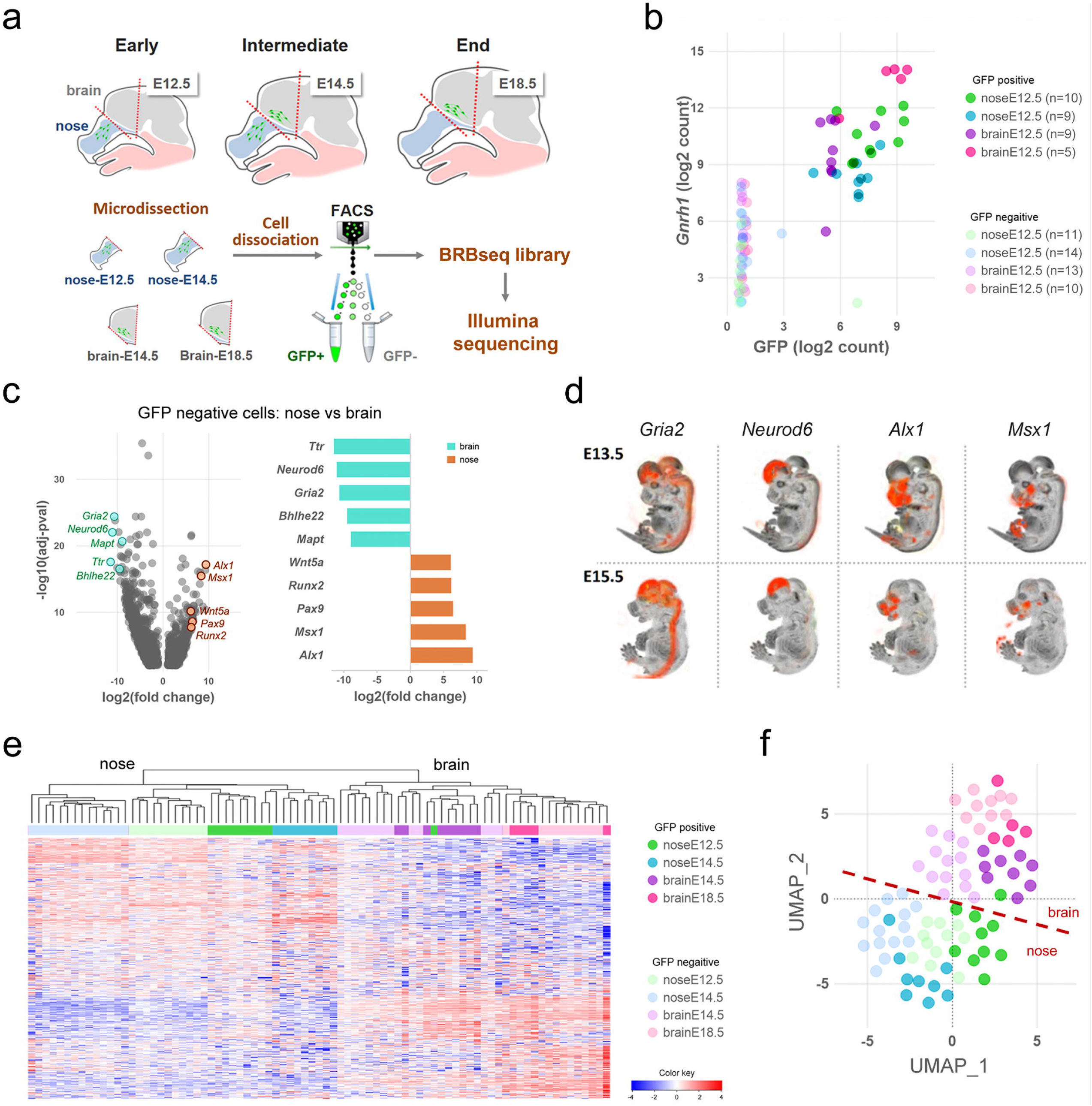
High resolution profiling of GnRH neurons in the mouse embryo. a) Strategy for isolation and RNA sequencing of GnRH+ and GnRH-cells neurons at different developmental stages. Nasal and brain region are microdissected from *GnRH::Gfp* mouse embryos at different developmental stages to follow GnRH neuron migration: early (E12.5 nose), intermediate (E14.5 nose), intermediate (E14.5 brain) and at the end (E18.5 brain). After cell dissociation and FAC isolation, GFP positive and GFP negative cells are processed for BRB library preparation followed by illumina sequencing. b) Expression analysis of Gnrh1 relative to Gfp validate the enrichment of GnRH positive neurons in the GFP+ fraction. c) Differential gene expression analysis of GFP negative cell comparing all nasal (orange) versus all brain (cyan) samples highlights specific regional markers. d) *In situ* hybridization expression levels of nose and brain

Surprisingly, cell localization (i.e., nose vs. brain) was the main driver of the first branching in the sample tree and the primary source of variation using hierarchical clustering on the 1000 most variable genes. Cell type (i.e., GnRH^+^ vs. GnRH^-^ cells) occurred at a lower hierarchical level (Fig. 1e). Consistently, dimensional reduction^18^ and principal component analysis (PCA) confirmed that nose and brain GnRH neurons clustered in two sub-populations (Fig. 1f, Supplementary. Fig 1).

### Molecular identity of GnRH neurons

The current knowledge of GnRH neuron molecular features is limited, with *Gnrh1* being the only marker of this neuronal population.^19^ We used Spearman’s correlation coefficient to rank all genes according to their correlation with Gnrh1 across the entire dataset to gain deeper insights into their molecular identity. Among the top 50 correlated genes (Fig. 2a, Supplementary Table 1), we identified transcription factors essential for GnRH expression, such as *Dlx5* and *Six6.* ^20-22^ *In situ* hybridization of these genes revealed expression profiles following the expected GnRH pattern in the mouse embryo, with some staining in other structures, including the olfactory epithelia and the forebrain (Fig. 2b). Notably, the top-ranked gene on our list was *Isl1*, a transcription factor recently identified in developing GnRH neurons in mice and humans.^23^

**Figure 2.**
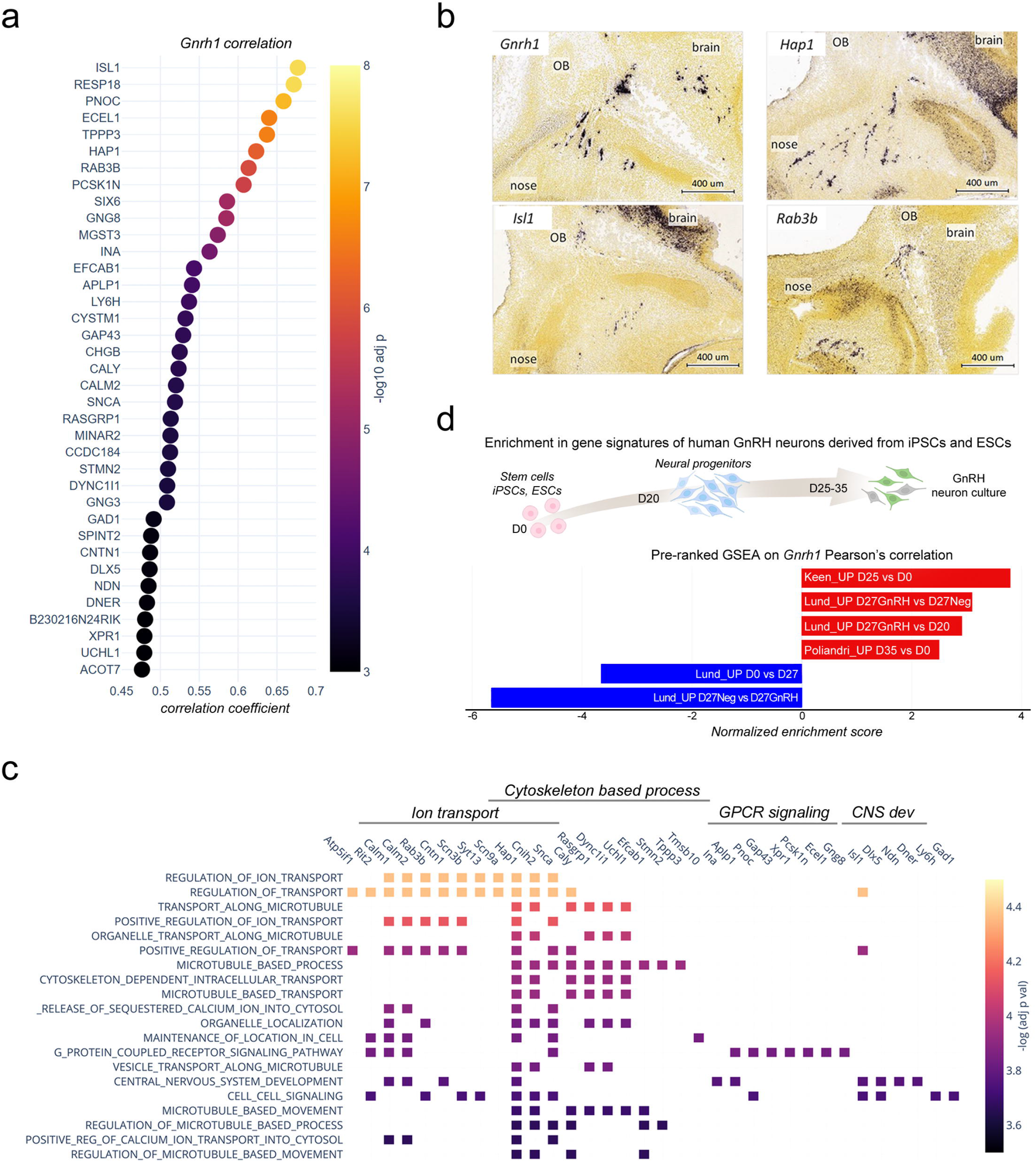
GnRH neuron molecular identity. a) Top 50 genes correlating with *Gnrh1* ranked by Spearman’s correlation coefficient. b) Representative *In situ* hybridization of *Gnrh1* and top correlating genes illustrating their similar expression patterns in sagittal sections from mouse embryos at intermediate developmental stages (E13.5-E15.5). OB, olfactory bulb. Data source: Allen Brain Atlas. c) Gene sets overlap of the top 50 signature genes with biological functions from the MSigDB database. CNS dev, central nervous system development. d) Gene set enrichment analysis (GSEA) with custom gene sets derived from RNAseq data of GnRH neuron cultures derived from iPSCs: Keen_UP D25 vs D0 (genes upregulated in GnRH neuron cultures after 25 days of differentiation *in vitro* vs. iPSCs from *Keen et al 2021*); Lund_UP D27GnRH vs D27Neg (genes upregulated in GnRH neuron cultures after 27 days of differentiation *in vitro* vs GnHR negative cells from *Lund et al., 2020*); Lund_UP D27GnRH vs D27Neg (genes upregulated in GnRH neuron cultures after 27 days of differentiation *in vitro* vs immature cultures at day 20 from *Lund et al., 2020*); Poliandri_UP D35 vs D0 (genes upregulated in GnRH neuron cultures after 35 days of differentiation *in vitro* vs. ESC from *Poliandri et al., 2017*).

Further, most top-ranked genes correlating with *Gnrh1* overlap with functional gene sets associated with microtubule-based function, ion transport, and GPCR signaling (Fig. 2c). *Stmn2* and *Stmn3* encode for Stathmins, microtubules-destabilizing proteins previously implicated with GnRH neuron migration.^24^ *Gap43*, encoding for the growth-associated protein 43, is expressed in developing olfactory and GnRH systems, although its biological role is not yet elucidated.^25,26^ *Dync1i1* encodes for a subunit of a microtubule-based motor protein, which is involved with split-hand-foot-malformation.^27,28^ a known CHH-associated phenotype.^29^ The genes involved in ion transport encode voltage-gated sodium channels (*Snc3b* and *Scn9a*), but also proteins modulating the activity of voltage-gated calcium channels (*Cbarp*) or controlling ion channel localizations (*Hap1*).

One central question related to the molecular identity of GnRH neurons is its conservation across species. To investigate the mouse-human similarity, we used available RNAseq data from recent studies describing the *in vitro* differentiation of GnRH neurons from human ESCs and iPSCs.^30-32^ Indeed, gene set enrichment analysis revealed a positive association of the mouse genes correlating with *Gnrh1* with gene signatures of human GnRH neurons. (Fig. 2d).

Altogether, these findings indicate that the molecular identity of GnRH neurons is complex and may involve a diverse set of transcription factors and other regulatory genes beyond *Gnrh1*.

### Spatiotemporal expression trajectories define GnRH neuron developmental stages

Using transcriptome analysis of GnRH-positive neurons, we identified wide transcriptional dynamics across different developmental stages (Fig. 3a, c). We employed a step-wise differential expression analysis across consecutive developmental stages to simplify this complexity by classifying genes into spatiotemporal trajectories (Supplementary Fig. 2, Supplementary Table 2). These trajectories parallel known biological processes occurring in GnRH neurons across development, such as the timely increase of N^12^ecdin and DLX transcriptional activators and decrease of MSX repressors modulating *Gnrh1* expression^20^ (Fig. 3b). Other examples are the "early stages trajectories" of genes encoding for syndecans, glypicans, and heparin sulfate proteoglycans which promote GnRH neuron migration ^12,33-35^ and the "brain trajectories" of genes involved in late stages, including genes encoding GABA receptor subunits^36,37^ and proteins critical for cell-cell adhesion such as PSA-NCAM, SynCAM and contactin^38-40^ (Fig. 3b).

**Figure 3.**
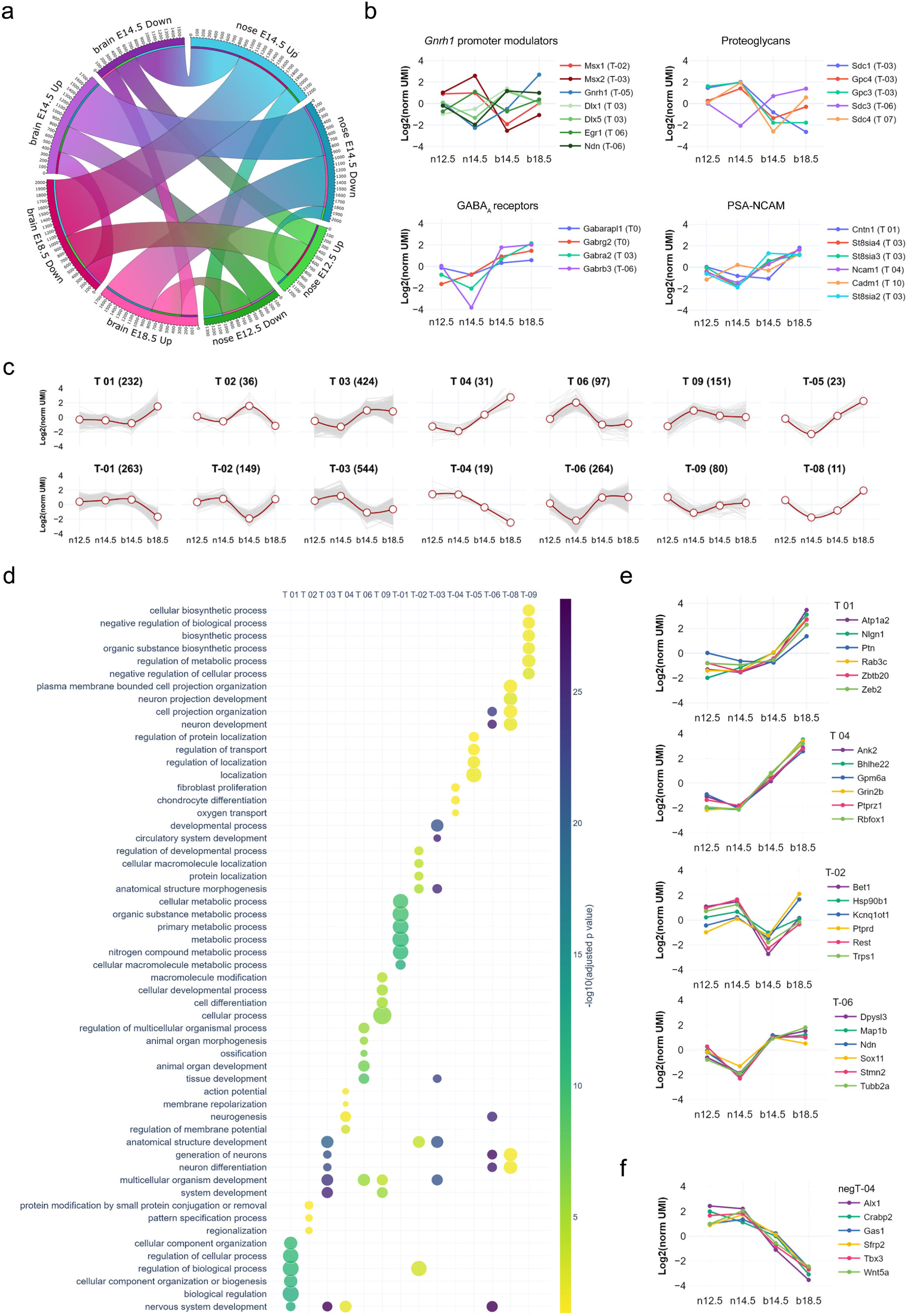
Spatio-temporal trajectories delineate GnRH neuron expression dynamics. a) Chord diagram representing the number of up- or down-regulated genes (arc thickness) after differential gene expression analysis of GnRH neurons across all developmental stages (colored nodes). b) Normalized expression profiles and trajectory classification of known genes involved in GnRH neuron development including modulators of *Gnrh1* promoter, proteoglycans, GABA_A_ receptors, and genes involved in the formation of Polysialic Acid Neural Cell Adhesion Molecule (PSA-NCAM). c) Expression dynamics of the main GnRH neuron trajectories (i.e. at least 10 genes per trajectory). Average gene expression is shown in red while grey lines represent individual genes. d) Scatter plot showing the top six significant terms in the main trajectories after functional enrichment analysis within biological processes from gene ontology database. e,f) Normalized expression profiles and trajectory classification of top genes emerging from representative trajectories significant after functional enrichment analysis.

Next, we combined spatiotemporal trajectories with functional enrichment analysis to identify spatially and temporally restricted biological processes throughout GnRH neuron development. Distinct biological processes were associated with different trajectories with slight overlaps (Fig. 3c, d and Supplementary Fig. 4), consistent with the idea that specific gene trajectories may drive spatiotemporally restricted biological functions to shape GnRH neuron development. Next, we selected the most differentially expressed genes associated with previously identified top GO terms (Fig 3e, Supplementary Fig. 5). Some genes were already linked with GnRH neuron biology or reproductive phenotype. Indeed, consistent with their trajectories, the microtubule-associated protein stathmin (*Stmn2*) and the repressor element-1 silencing transcription factor (*Rest*) promote migration in immortalized GnRH neurons.^24,41^

The trajectory analysis of GnRH-negative cells revealed T-box transcription factor 3 (*Tbx3)* as a gene with high expression in the early stages in the nasal compartment (Fig. 3f). Notably, its human orthologue is mutated in patients with ulnary mammary syndrome (OMIM# 181450), a condition associated with CHH.^42^ Another candidate in the same trajectory is a member of the Wnt family (*Wnt5a*). This expression profile is consistent with previous detection in the olfactory mesenchyme^43^ and its ability to activate olfactory ensheathing cells,^44^ a population of migrating glial cells providing a permissive microenvironment and guidance for GnRH neuron migration.^45^

Candidates upregulated in GnRH neurons at the end of migration are of particular interest. We found *Nlgn1* (T 01), which belongs to the neurexins and neuroligins family, a group of cell-cell adhesion molecules involved in synaptic formation and plasticity.^46^ Notably, *Nlgn3,* another member of this family, has a similar trajectory (T 03), and mutations in its human orthologue have been identified in CHH patients.^47^ Another appealing candidate with increased expression at the end of migration (T 01) is Pleiotrophin (*Ptn*), a secreted heparin-binding growth factor that binds to the perineuronal net (PPNs).^48^ Pleiotrophin binds with high-affinity Posphacan (*Ptprz1*) and has been shown to modulate migration and induce neurite outgrowth^48-50^ (Fig. 3e). Notably, several *Ptprz1* paralogs (*Ptprs*, *Ptprd*, *Ptpro*, *Ptprf*) are dynamically expressed in GnRH neurons and, together with the other member of their family, have been implicated with the timing of puberty through genome-wide association studies (GWAS).^51^

Altogether, these findings show that complex expression dynamics are a distinctive feature of genes controlling GnRH neuron development. These genes drive the timely activation of key biological processes and can be identified by spatiotemporal trajectory analysis.

### Dynamic networks of cell-to-cell communication highlight timely signaling pathways for GnRH neuron development

Based on the extent of the observed transcriptional dynamics, we anticipated that the cell-to-cell-communication (CCC) between GnRH neurons and their surrounding environment could be affected by changes in specific ligands (environment) and receptors (GnRH neurons). CCC analysis has recently improved thanks to the growing availability of accurate protein-protein interaction (PPI) databases and is now widely used to construct tissue and cell-specific ligand-receptor networks by inferring active signaling pathways based on synchronized expression from transcriptomic data.^52^

We combined a curated database of ligand-receptor interactions (CellTalkDB)^53^ with our trajectory analyses on GnRH neurons and GnRH-negative cells to construct a dynamic protein-protein interaction network of directional paracrine communication between the environment and GnRH neurons. Gene trajectories were annotated on ligand and receptor nodes, while the strength of ligand-receptor interactions was calculated based on the expression correlation of ligands and receptors in each pair (Fig. 4a).

**Figure 4.**
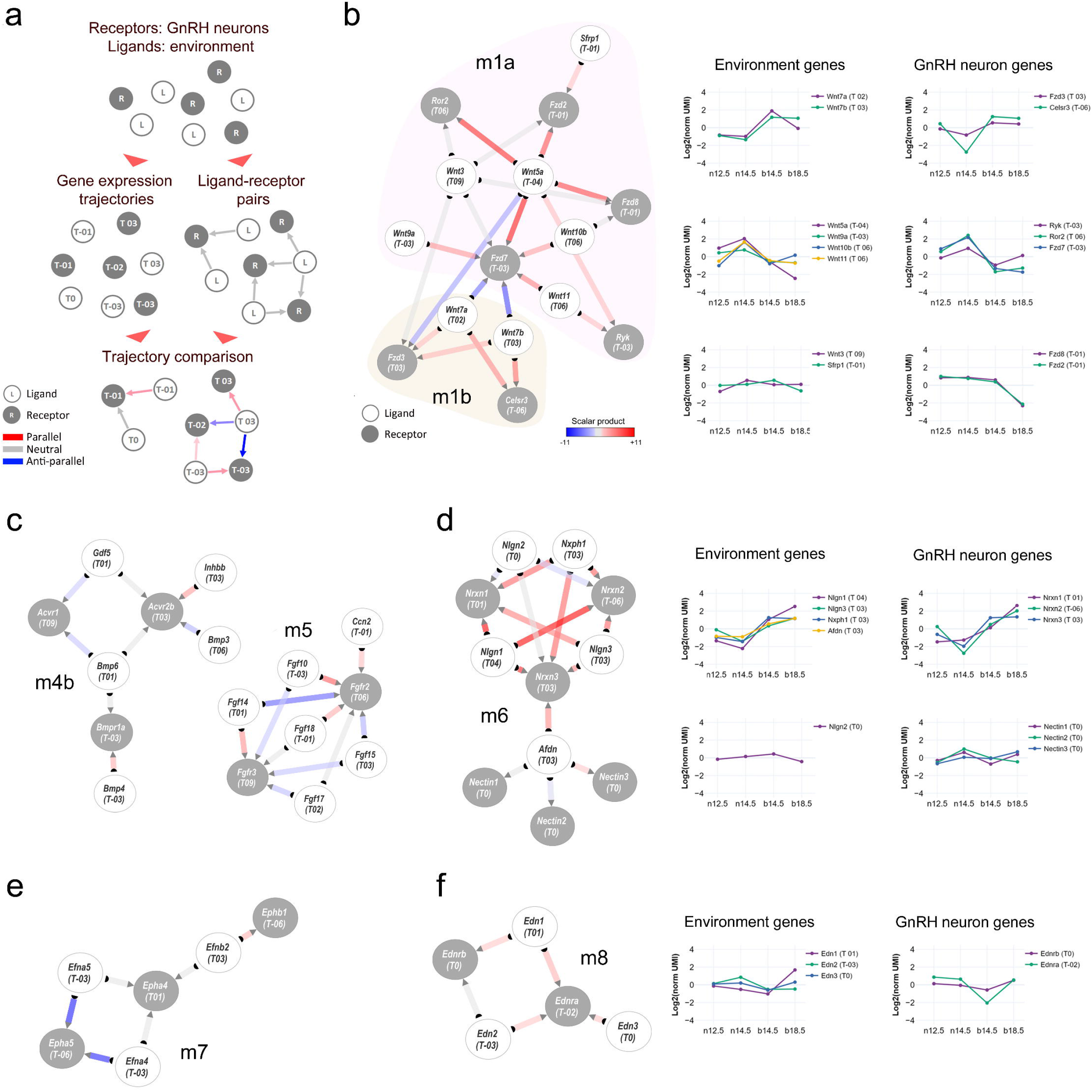
Dynamic cell-to-cell communication networks in GnRH neuron development. a) Schematic illustrating the strategy to obtained dynamic interactions between GnRH neurons and their environment by combining PPI data with gene expression trajectories (see methods). PPI network and gene expression profiles of different modules of ligands and their respective receptors including b) Wnt submodules m1a and m1b, c) BMP submodules m4b and FGF module m5, d) Neurexins and Neuroligins in module m6, e) ephrins module m7, and f) endothelin module m8. Node color discriminate between receptors expressed in GnRH neurons (grey) and ligands expressed in the environment (white). The degree of coordinated expression (i.e., scalar product) is color coded on edges.

Based on the local connectivity and the inferred strength of ligand-receptor pairs, we used the resulting network to identify different functional modules and sub-modules. As expected, many of those contained gene families and pathways already linked with GnRH neuron development, such as semaphorin family members and their receptors, plexins, and neuropilins (Supplementary Fig. 6, module m3). Semaphorins play a crucial role in brain development and GnRH neuron development, with different members of this signaling pathway eliciting opposing effects depending on the molecular environment.^54-56^ Other modules display elements of the Wnt (m1), BMP (m4b), and FGF (m5) signaling pathways (Fig. 4b, c), which are well known for their cooperative role in the patterning of the olfactory placode and its neurogenic niche.^57^ Consistent with these functions, we found one of the two Wnt submodules (m1a: *Wnt5a, Wnt9a, Wnt10b, Wnt11, Ryk, Ror2, Fzd7*) with an early stages activation profile. However, the second submodule has an opposite spatiotemporal profile (m1b: *Wnt7a, Wnt7b, Fzd3, Celsr3*), suggesting that some components of Wnt signaling might be involved in later events, including GnRH neuron maturation and connectivity. Our analysis highlights pathways previously linked with mouse GnRH neuron development in mice and humans like DCC/Netrin (m9),^58-60^ and strengthen old candidates such as ephrin (m7)^61^ and endothelin (m8)^62,63^ signaling pathways (Fig. 4e-f).

One of the most compelling findings of this analysis is the identification of the neurexins/neuroligins module (m6) presenting with highly coordinated expression at the end of the migratory phase (Fig. 4d). Neurexins are presynaptic transmembrane proteins that interact with postsynaptic neuroligins to form trans-synaptic complexes, which play a critical role in regulating synaptic function, and facilitating synapse formation and maintenance and could be involved in the establishment of GnRH neuron connections with other neurons.^46,64,65^

### GnRH neuron trajectories are linked with the genetics of human reproduction

Combining spatiotemporal trajectories with PPI, we identified genes critical for GnRH neuron biology in rodents and genes previously linked with human reproduction, including *NLGN3*, *DCC*, *NTN1*, and members of the PTPR, Semaphorin/Plexin, and Neuropilin families. These findings prompted us to investigate whether expression trajectories could reveal genetic association with relevant aspects of human reproduction in the general population and in patients with congenital GnRH deficiency (CHH).

We first used GWAS catalogue to retrieve a list of genes associated with human traits linked with reproductive onset (i.e., age at menarche, EFO_0004703; age at menopause, EFO_0004704; age at the first sexual intercourse, EFO_0009749). (Supplementary Table 4). We used these genes to test their enrichment against the GnRH neuron trajectories (Fig. 5a) and found T01 as the top-ranked trajectory (P_adj_=5.8×10^-5^) followed by T03 (P_adj_=2.1×10^-2^). T01 trajectory contains genes upregulated at the end of GnRH neuron migration and controlling the formation of neuronal projections and synaptic contacts (Supplementary Table 2).

**Figure 5.**
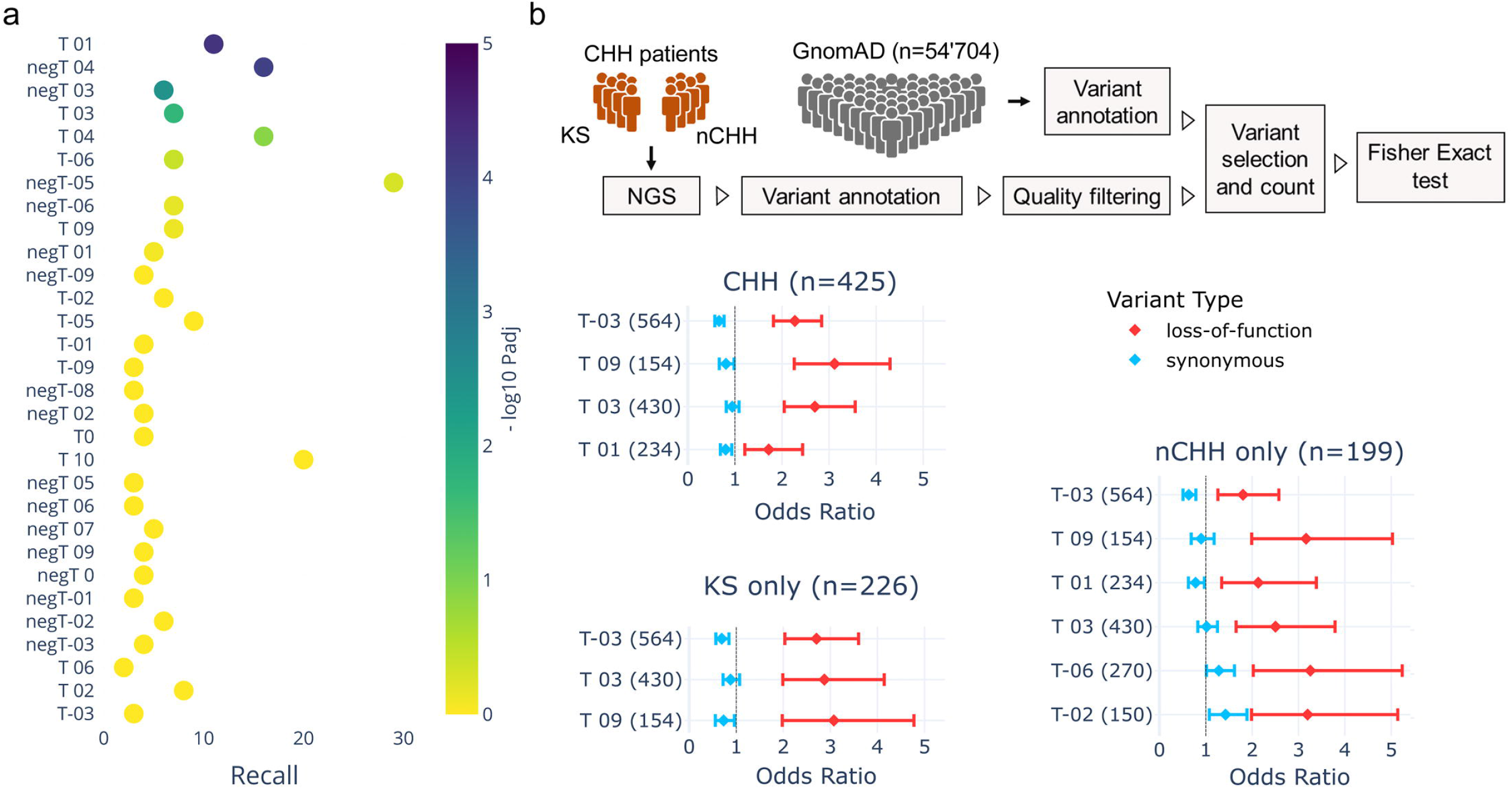
Genetic overlaps linking GnRH neuron trajectories with human reproduction. a) Scatter plot illustrating the enrichment across GnRH neuron trajectories of GWAS genes associated with reproductive onset (i.e., puberty, age at menarche, age at first sexual intercourse, and age at menopause from GWAS catalog). Trajectories below the threshold of significance are colored in yellow. b) Forest plots showing the mutation load of rare PTVs relative to synonymous in patients with CHH, KS, and nCHH vs GnomAD controls. Error bars represent the 95% confidence interval of the odds ratio.

We then used a large cohort of CHH patients (n=425) and GnomAD controls to perform a trajectory-collapsed burden test of rare protein-truncating variants (PTVs) relative to synonymous (Supplementary Table 5). Four trajectories showed significant enrichment: T-03 (P_adj_=2.3×10^-18^), T09 (P_adj_=1.9×10^-11^), T03 (P_adj_=3.7×10^-10^), and T01 (P_adj_=1.0×10^-3^) (Fig. 5b). To get a complete picture of the involvement of these trajectories, we repeated the analysis taking into account the KS and normosmic CHH (nCHH) sub-phenotypes, which are well known to display different genetic architectures associated with distinct pathogenic mechanisms.^66,67^ Consistently, the enrichment of some trajectories was mainly driven by PTVs in KS patients such as T-03 (KS P_adj_=2.1×10^-13^ nCHH P_adj_=10^-5^) and T03 (KS P_adj_=2.9×10^-7^ vs. nCHH P_adj_=1.6×10^-3^), (Fig. 5b Supplementary Table 6 and 7). In contrast, T01 (P_adj_=1.3×10^-3^), T-06 (P_adj_=6.1×10^-3^), and T-02 (P_adj_=4.6×10^-2^) were exclusive to nCHH.

Altogether, these results show that GnRH neuron trajectories are linked with genetic determinants driving both physiological and pathological variability of human reproduction.

## Discussion

Our study provides high-resolution transcriptomic profiling of GnRH neurons during migration in the mouse embryo. Analyzing more than 20’000 GnRH^+^ cells from the mouse embryo, this work delineates the rich expression dynamics of GnRH neurons along four pseudo-timepoints. This work expands the scope of previous studies^68-71^ and enhances our understanding of the molecular mechanisms underlying GnRH neuron ontogeny.

We identified a deep transcriptional shift from the nose to the brain when GnRH cells drastically change their environment. This shift transcends the variability between cell types (GnRH^+^ and GnRH^-^ cells) and involves the upregulation of genes shaping neuronal activity (e.g., *Snc3b* and *Scn9a,* encoding voltage-gated sodium channels; *Grina* and *Grin2b* encoding subunits of the glutamate ionotropic receptor). This is reminiscent of a recent study in zebrafish larvae showing a pause of the migration of the GnRH cells at the nasal-forebrain junction (NFJ), during which they acquire coordinated neuronal activity required to enter the brain.^72^ Notably, GnRH neurons have been identified as clusters of cells accumulated in this region (cribriform plate in humans) instead of entering the brain in a human KS fetus.^3^

One conundrum in the study of the biology of GnRH neurons is the reliance on *Gnrh1* as the sole marker. Our analysis identified several high-ranked genes correlating with *Gnrh1* across development, which could help define GnRH neuron identity. Among these genes, *Isl1,* the top ranked-gene, plays a crucial role in the migration, axonal pathfinding, and maturation of peripheral neurons.^73-75^ *Isl1* has been proposed as a candidate gene for GnRH neuron development^23^ and is also enriched in cultures of GnRH neurons derived from human induced pluripotent stem cells (iPSC).^31^ Genes highly co-expressed with *Gnrh1* appear to be involved in specific biological processes although not unique to GnRH neurons (i.e., cytoskeleton remodeling and ion transport). Further studies using single-cell analysis are needed to assess which combinations of genes better characterize GnRH neurons’ molecular identity across development.

Resolving gene expression dynamics based on spatiotemporal trajectories in GnRH^+^ and GnRH^-^ cells highlights specific biological processes at different developmental stages. This is exemplified in the timely increase of Necdin and DLX transcriptional activators and decrease of MSX repressors promoting the progressive rise of *Gnrh1* promoter activity. We noticed a slight overlap between biological processes associated with specific trajectories consistent with the idea that each trajectory may shape biological networks and signaling pathways for GnRH neuron development (T01 neuronal maturation and connectivity). Combining these expression trajectories with PPI networks allowed us to infer cell-to-cell communication dynamics supported by the identification of known signaling modules (e.g., Semaforins, DCC). Further, this analysis unravels novel candidate gene pathways such as Neurexins and Endothelins.

Finally, our data revealed an emerging link between trajectories and genetic basis of human reproduction. Previous GWAS studies identified many loci associated with the reproductive traits in humans,^76-78^ including genes affecting fertility via energy homeostasis or directly controlling the activity of GnRH neurons or their activators (i.e., Kisspeptin neurons).^79-81^ Our study uncovers a novel link between reproductive onset and the late embryonic development of GnRH neurons by showing a specific enrichment genes in T01 and T03 trajectories. Consistently, these trajectories contain genes involved in establishing neuronal projections and synaptic contacts, which likely help integrate GnRH neurons into the hypothalamic network that controls puberty. Notably we have successfully replicated these results in another study using 660 age at menarche genes identified after genetic analysis in ∼800,000 women from UK Biobank. ^82^

In contrast, genes involved in congenital GnRH deficiency have already been linked with biological events critical for GnRH neuron development.^66,83-85^ In line with this, we measured a genetic burden of PTVs (CHH patients *vs*. controls) in trajectories mirroring the shift of GnRH neurons from nose to brain. Accordingly, when we focused on KS sub-phenotype, we found a more robust enrichment for T03 T-03 trajectories. Further, we encounter a specific association between nCHH and the trajectory T01 which we also linked with reproductive onset.

In conclusion, the comprehensive analysis of gene expression dynamics in GnRH neurons during embryonic development expands our understanding of the underlying molecular mechanisms. We highlighted the importance of GnRH neuron expression trajectories in coordinating crucial biological processes across embryonic development and their links with critical aspects of human reproduction, such as the reproductive onset and infertility.

## Methods

### Animals

*Gnrh::Gfp* mice, a generous gift of Dr. Daniel J. Spergel (Section of Endocrinology, Department of Medicine, University of Chicago, IL),^16^ were housed at room temperature (22°C) with a 12-h-light/12-h-dark cycle and free access to water and food. All experimental protocols were performed following the Swiss animal welfare laws under the authorization of the Service de la consummation et des affaires vétérinaire Vaud. *Gnrh::Gfp* embryos were harvested at E12.5, E14.5, and E18.5 (plug day, E0.5) and used for the microdissection of nasal and brain regions for GnRH neuron isolation.

### Isolation of GnRH neurons using fluorescence-activated cell sorting

The nose and forebrain microdissections from *Gnrh::Gfp* embryos were enzymatically dissociated using a Papain Dissociation System (Worthington, Lakewood, NJ) to obtain single-cell suspensions.^68,86^ Cells were stained with a far-red cell membrane-permeant nuclear dye (RedDot™1, Biotium) to specifically select live cells and DAPI to exclude dead cells. Immediately after staining, FACS was performed on a MOFLO ASTRIOS EQ (BD Bioscience), and, for each microdissection, 500 to 1500 GFP+ and 2000 GFP-cells were sorted directly into 10 μl of extraction buffer.

### RNA processing and transcriptomic profiling

Extraction of total RNA was carried out using Acturus PicoPure™ RNA Isolation Kit (Thermofisher) following the manufacturer’s protocol. Concentration and integrity were assessed using the Qubit RNA HS Assay Kit (Life Technologies) and High Sensitivity RNA ScreenTape Assay (Agilent), respectively. Libraries were prepared following the BRB-seq protocol using at least 15 ng total RNA per sample. Sequencing was performed using the Illumina NextSeq 500 platform, and data preprocessing, including sample demultiplexing and alignment, was performed to obtain gene count matrices.^87^

### Differential gene expression, functional classification, and enrichment analysis

We performed gene expression analysis on RNAseq data using the edgeR quasi-likelihood method.^88^ Raw count data were first preprocessed and normalized for differences in library size and composition. Differential expression analysis was then performed using a quasi-likelihood framework, which models both biological variability and technical noise. If not otherwise indicated, genes were considered differentially expressed if they had a false discovery rate (FDR) adjusted p-value below 0.05 and a fold change greater than 2.

Spatiotemporal trajectory classification was performed separately on data for GnRH neurons and GFP^-^ cells. First, differential gene expression analysis was performed in each of the three transitions between the four developmental steps. Then, all transitions were classified according to statistical significance (pval<0.01; fold change > 2), and each gene was annotated with an array combining the results of the three consecutive transitions (Supplementary. Fig 2 and 3).

Computation of gene set overlap for GnRH neuron signature genes has been performed using MSigDB resources (http://www.gsea-msigdb.org/gsea/msigdb/annotate.jsp).^89-91^ Gene set enrichment analysis (GSEA) was used with custom gene sets derived from transcriptomic data of GnRH neuron cultures derived from induced pluripotent stem cells (iPSCs).^30-32^ The significance of enrichment was evaluated by calculating a normalized enrichment score (NES), and gene sets with an FDR-adjusted p-value < 0.05 were considered significantly enriched. Finally, we used g:GOSt web server (https://biit.cs.ut.ee/gprofiler/gost)^92^ for functional enrichment analysis with default and custom datasets.

### Dynamic Cell-to-cell communication network

For cell-to-cell communication analysis, we used CellTalkDB (http://tcm.zju.edu.cn/celltalkdb),^53^ a manually curated database of ligand-receptor pairs, to annotate all receptors expressed in GnRH neurons as well as ligand expressed in the environment, and build a ligand-receptor interaction network (Fig. 4a). Next, we classified both ligand and receptor genes according to the corresponding trajectories (Supplementary Fig. 2 and 3). Finally, the strength of ligand-receptor interactions was inferred by the expression profiles of ligand-receptor pairs using the scalar product of their normalized expression vectors. We obtained an interaction network representing the dynamic interaction between GnRH neurons and their environment during embryonic development (Supplementary Fig. 4). Finally, functional modules and sub-modules were identified based on their local connectivity (i.e., number and strength of edges).

### Next-generation sequencing of CHH patients

The CHH cohort included 226 KS and 199 normosmic CHH (nCHH) for a total of 425 unrelated probands (320 males and 105 females) whit a majority of European descent. All subjects provided written informed consent, and their clinical phenotype was assessed as previously described.^93^

After DNA extraction, paired-end whole exome sequencing was performed at the Denmark facility of BGI (Beijing Genomics Institute) Global (n=106) or at Health 2030 Genome Center (a portion of subjects between 2019 and 2021, n=124), while whole genome sequencing (n=195) was performed using DNBSEQ technology through the Denmark facility of BGI (Beijing Genomics Institute) Global.^94^ Briefly, the resulting raw sequences (fastq files) are processed by an in-house bioinformatics analysis workflow that relies on Sentieon DNASeq, a GATK-compliant toolbox that maps the reads to the human reference sequence (GRCh37) and detects variants.^95,96^ Identified variants are then annotated with minor allele frequencies (MAFs) from gnomAD (v2.1.1, n=54’704 controls) (http://gnomad.broadinstitute.org) and with multiple pathogenicity prediction tools^97,98^ using ANNOVAR.^99^ The GnomAD exomes vcf file was downloaded and annotated with ANNOVAR using the same databases to ensure coherence.

For the genetics burden, we applied a series of filters to minimize false positive calls and sequencing artifacts while preserving truly positive calls. This strategy, detailed below, was used for both the CHH cohort and the GnomAD controls whenever possible. We excluded variants with a popmax frequency in GnomAD higher than 0.01%, given the prevalence of CHH.^66^ We retained nonsense variants (i.e., stop gain, frameshift, acceptor-donor splice sites ± 2bp from an exon or SpliceAI^100^) that passed strict filters guided by GATK recommendations (minimum quality score of 50, mapping quality > 55 and Mapping Quality Rank Sum Test > -2.5) [https://gatk.broadinstitute.org/hc/en-us/articles/360035890471-Hard-filtering-germline-short-variants]. Variants with an allelic depth ratio under 20% or located in segmental duplications^101^ were discarded. Putative private or ultra-rare variants that were frequent (> 3 times) in a local genetic database of healthy individuals (n=300) were also removed, as they were considered systematic sequencing artifacts. Furthermore, variants that were present in excess of pedigrees (>5) or flagged in GnomAD, as well as indels involving more than 3 nucleotides, were excluded, due to the higher error rate in calls, primarily from alignment issues. After applying all the filters, the subjects with at least one variant in the same trajectory were counted. Each variant in GnomAD’s data was assumed to come from a different individual, with a ceiling at the number of controls, and a contingency table with affected and wild-type alleles was constructed. A one-sided Fisher’s Exact test for each trajectory was then performed to estimate the enrichment of variant alleles in the CHH cohort vs. GnomAD controls. This analysis was applied on synonymous variants through the same filters to control for overall inflation of numbers from potential unaccounted for sources.^102^ From the odds ratio obtained, a 95% confidence interval for the odds ratios could be estimated as follows:

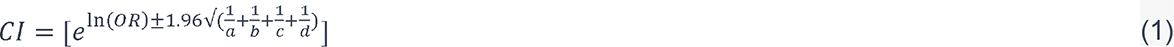

with a,b,c and d being the count values from the contingency table. Using (1), it is then possible to calculate a z-score statistic between the estimated odds ratios from PTVs and synonymous variants following:

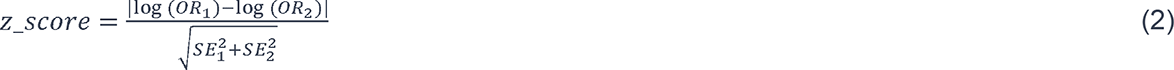

with 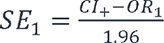. Only the trajectories passing a Bonferroni correction were retained and considered as having a significant enrichment of PTVs relative to synonymous (i.e., the loss-of-function burden test has a significantly higher odds ratio than the synonymous burden testing CHH). Finally, this process was repeated separately for nCHH and KS probands to evaluate phenotype specificity.

