## Supplementary Figures for "Transcriptomic profiling of murine GnRH neurons reveals developmental trajectories linked to human reproduction"

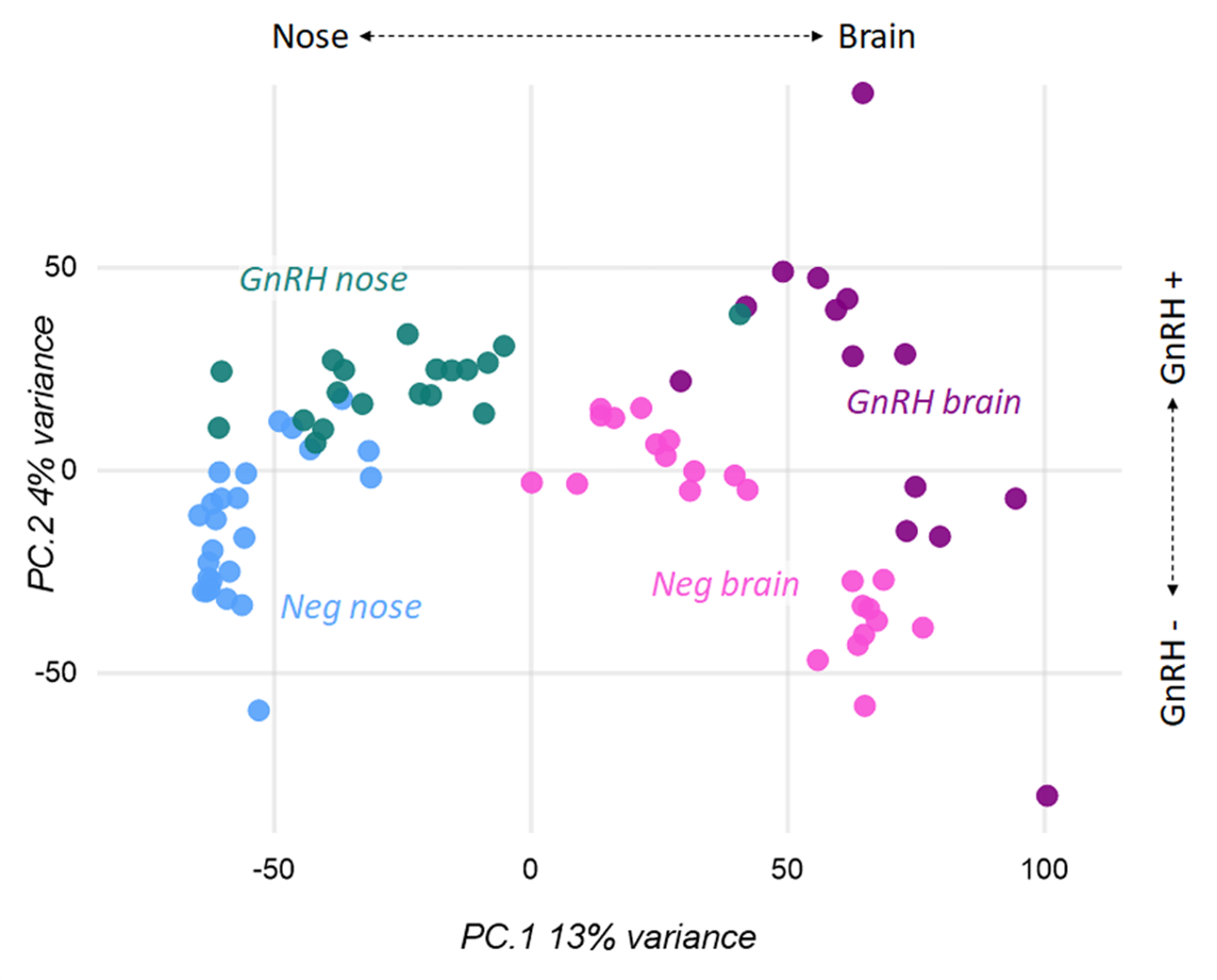

Supplementary Figure 1 | **PCA of RNA sequencing from GnRH^+^ and GnRH^-^ mouse embryonic cells.**

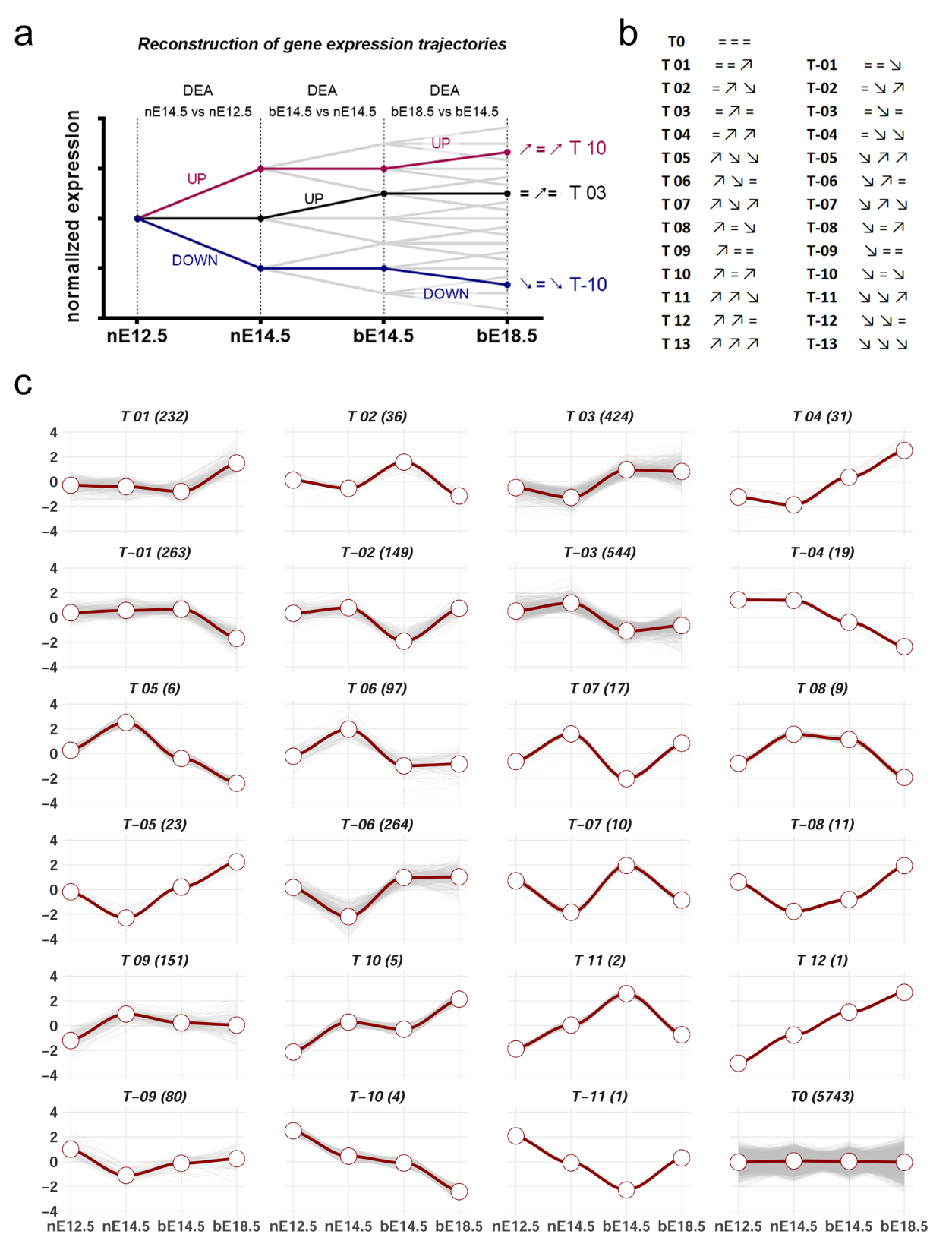

Supplementary Figure 2 | **GnRH^+^ cells gene expression dynamics**. a) Algorithm to assign trajectory names to expression dynamics. b) Pattern description of all possible trajectories in the GnRH transcriptomics dataset. c) Expression of all detected genes in GnRH+ cells organized by trajectory. Average gene expression is shown in red while grey lines represent individual genes.

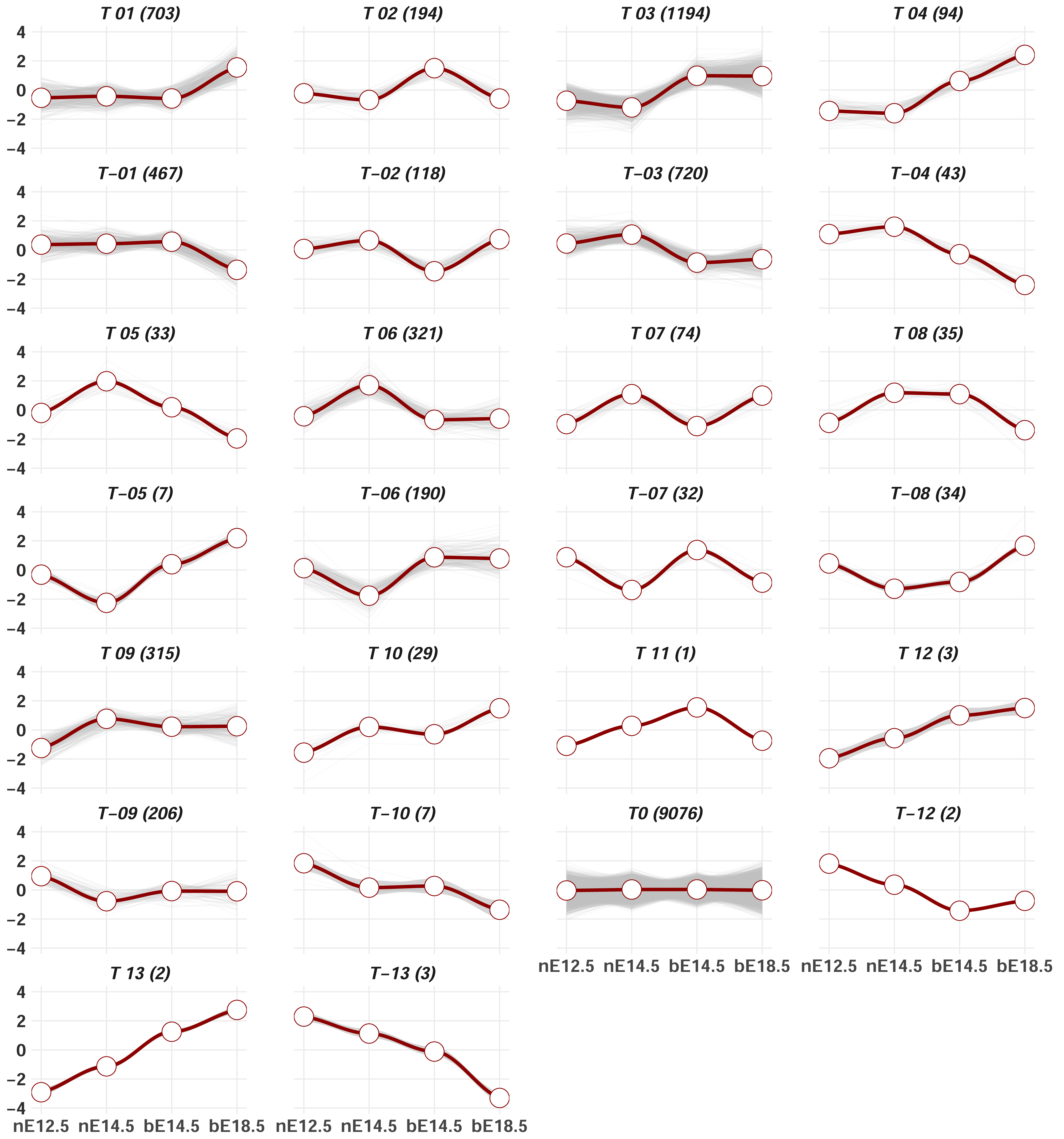

Supplementary Figure 3 | **GnRH^-^ cells gene expression dynamics**. Expression of all detected genes in GnRH- cells organized by trajectory. Average gene expression is shown in red while grey lines represent individual genes.

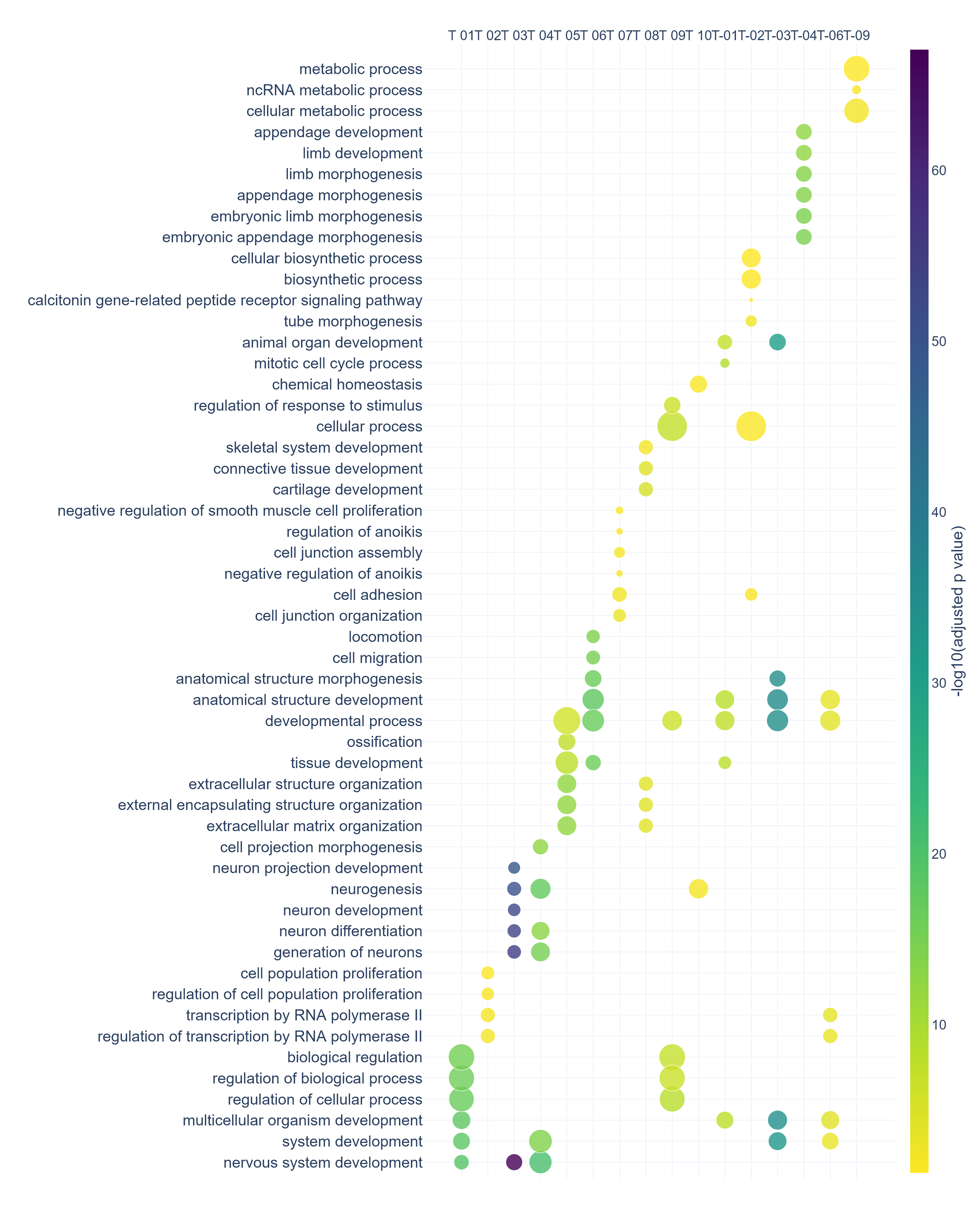

Supplementary Figure 4 | **Functional enrichment analysis in GnRH^-^ cells**. Scatter plot showing the top six significant terms in the main trajectories of GnRH- cells after functional enrichment analysis within biological processes from gene ontology database.

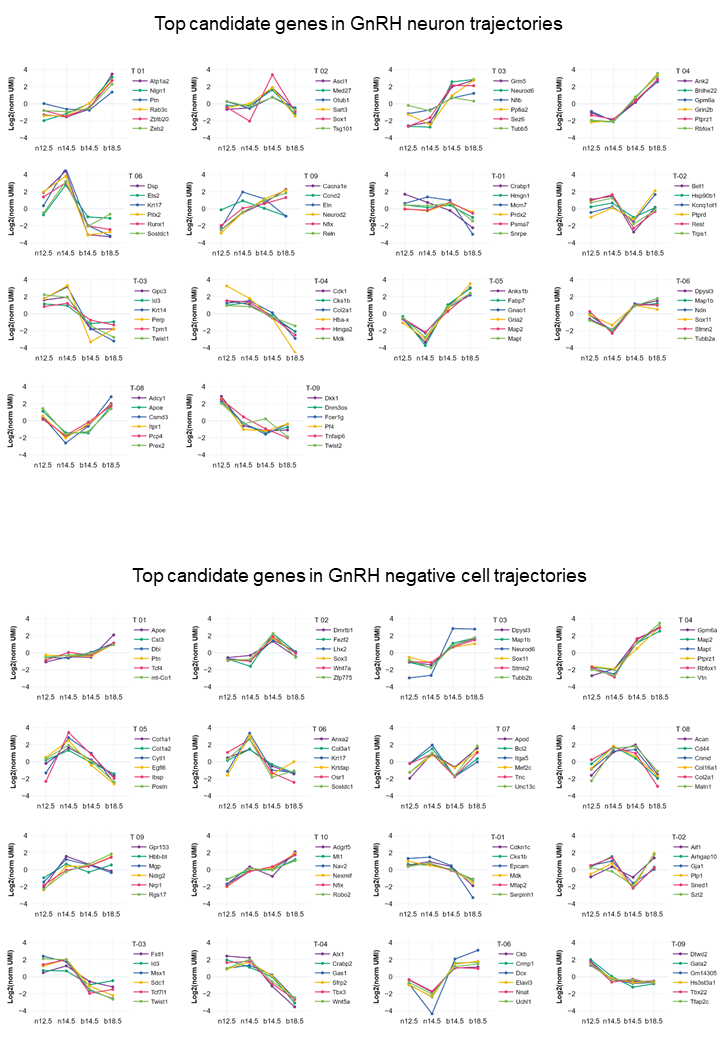

Supplementary Figure 5 | **Expression profiles of top genes from trajectories enriched in biological processes**. Normalized expression profiles and trajectory classification of top genes emerging from representative trajectories significant after functional enrichment analysis in both GnRH^+^ and GnRH^-^ cells.

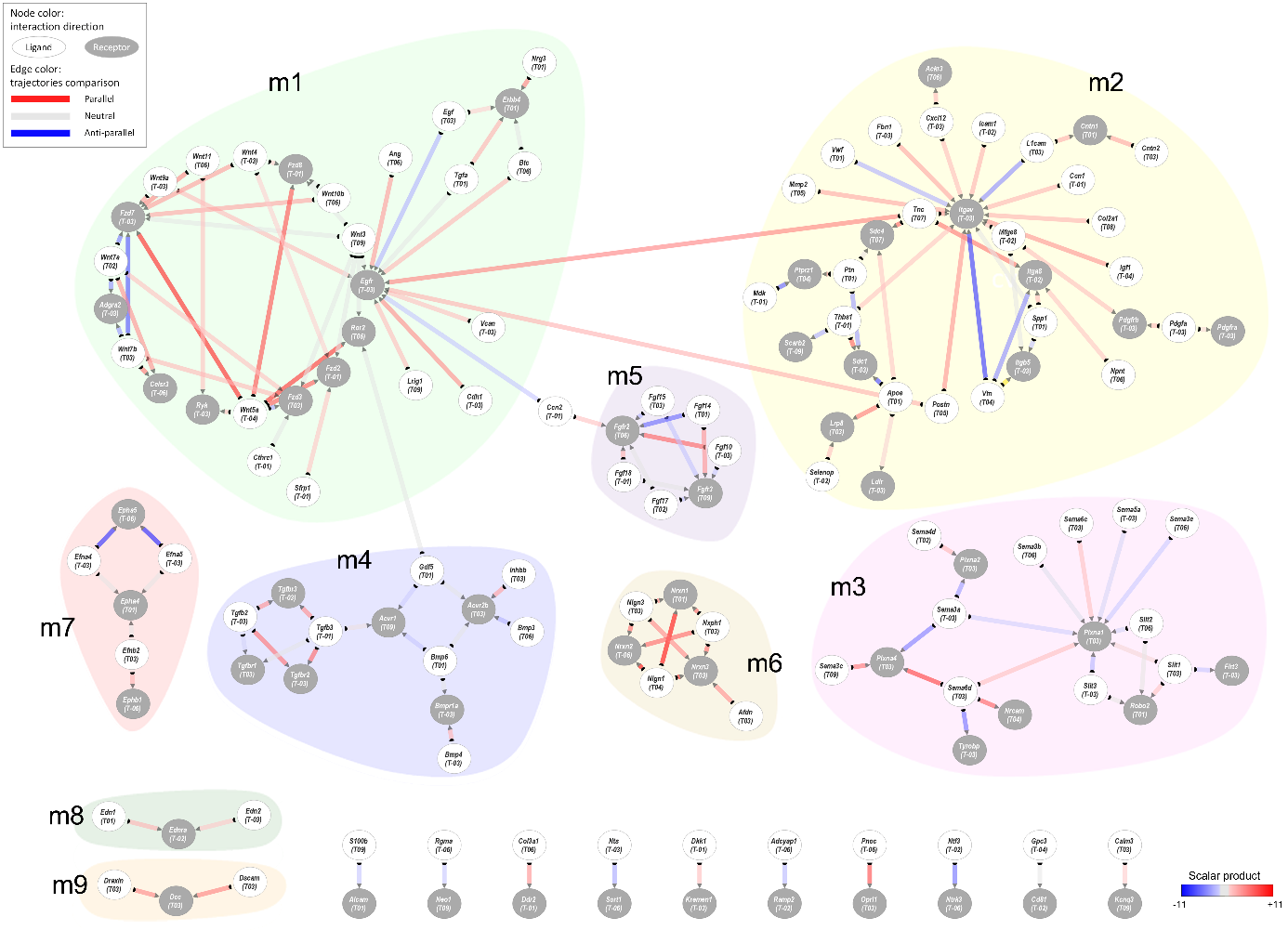

Supplementary Figure 6 | **Full dynamic cell-to-cell communication networks in GnRH neuron development.** PPI network and gene expression profiles of different modules of ligands and their respective receptors including (T0 nodes were hidden for visual clarity).
